# The extracellular matrix protein fibronectin modulates metanephric kidney development

**DOI:** 10.1101/2022.07.01.497442

**Authors:** Kathrin Skoczynski, Andre Kraus, Maike Büttner-Herold, Kerstin Amann, Mario Schiffer, Kristina Hermann, Leonie Herrnberger, Ernst R. Tamm, Bjoern Buchholz

## Abstract

Complex interactions of the branching ureteric bud and surrounding mesenchymal cells during metanephric kidney development determine the final number of nephrons. Alterations that result in impaired nephron endowment predispose to arterial hypertension and chronic kidney disease. In the kidney, extracellular matrix (ECM) proteins are usually regarded as acellular scaffolds or as the common histological end-point of chronic kidney diseases. Only little is known about their physiological role in kidney development. The ECM protein fibronectin is expressed at early time points of kidney growth. Therefore, we were interested in the characterization of its expression and role in kidney development. In mouse, fibronectin was expressed during all stages of kidney development with significant changes over time. At embryonic day (E) 12.0 and E13.5 fibronectin lined the ureteric bud epithelium, which was less pronounced at E16.5 and then switched to a glomerular expression in the postnatal and adult kidneys. Similar results were obtained in human kidneys. Deletion of fibronectin at E13.5 in cultured metanephric mouse kidneys resulted in reduced kidney sizes, impaired branching morphogenesis and decreased glomerulogenesis. In line with these findings, fibronectin deletion led to reduced cell proliferation of the branching epithelial cells. Fibronectin colocalized with alpha8 integrin and fibronectin loss caused a reduction in alpha8 integrin-dependent formation of glial cell line-derived factor and expression of Wnt11, both of which are essentially involved in promoting ureteric bud branching. In conclusion, the ECM protein fibronectin acts as a regulator of metanephric kidney development and is a determinant of nephron number.

## Introduction

Embryonic kidney development is a complex process, which may explain why urinary tract malformations are among the most common birth defects in humans.^1,2^ In humans, the development of the final kidney, the metanephros, starts at embryonic day (E) 35 with reciprocal signaling between the Wolffian duct and the adjacent metanephric mesenchyme (MM).^3^ Nephrogenesis significantly depends on the formation of the ureteric bud (UB) and the nephron endowment at the branching UB tips.^4^ In mouse, kidney development begins at E10.5 with the outgrowth of the UB from the Wolffian duct, which then invades the MM.^4^ The UB, which later becomes the collecting duct system, begins to branch dichotomously and induces the MM to condense and undergo mesenchymal-to-epithelial transformation.^5^ The condensed MM then forms renal vesicles, comma- and s-shaped bodies, which later turn into the nephrons.^5^ The inductive signaling is an essential step in kidney development because it induces UB outgrowth and subsequent UB branching. The glial cell line-derived factor (GDNF) is produced by the MM cells and plays a key role in UB branching.^6^ GDNF promotes UB cell proliferation and regulates the expression of the extracellular signaling molecule WNT11 in the UB tips, which in turn increases GDNF expression in a positive feedback loop.^7^ GDNF deletion in mice results in significant defects in UB outgrowth and consequently renal agenesis.^6^

Integrins are cell adhesion receptors which interact with several proteins of the extracellular matrix (ECM) and are involved in embryonic kidney development.^8^ Integrins consist of the subunits α and β and are divided into three groups called laminin-, collagen- and RGD-binding integrins.^8^ The RGD-binding integrins are linked to several ECM ligands.^8^ The RGD-binding integrin α8β1 (ITGA8) is expressed by MM cells and stimulation of ITGA8 significantly contributes to GDNF induction.^9^ Similar to GDNF deletion, loss of ITGA8 leads to impaired UB outgrowth in the MM and results in renal agenesis.^6,9^

Next to the dramatic phenotypes provided by loss of GDNF or ITGA8, alterations in kidney development do not always have to result in renal agenesis but can affect the final kidney size and number of nephrons and glomeruli. In humans, nephron number between individuals varies from roughly 200,000 up to 2,000,000 nephrons, which exceeds the structural variability of most other organs.^10^ Low nephron number increases the risk for chronic kidney disease (CKD) due to lower functional reserve and continuous hyperfiltration.^11^ In addition, low nephron number also predisposes to arterial hypertension.^12,13^ Despite the significant impact of nephron endowment for health, the determinants of nephron number remain incompletely understood.

ECM proteins such as collagens, fibronectin, and laminins are often regarded as scaffolding proteins in the kidney providing structural stability or as an irrefutable histological sign of progressive CKD.^14^ However, ECM proteins are largely involved in signal transduction and therefore may affect kidney development as recently shown for the ECM protein nephronectin.^15^ Recently, fibronectin has been shown to mediate branching morphogenesis of mouse embryonic salivary glands and additionally was detected at early stages of mouse metanephric kidney development.^16^ Fibronectin is a glycoprotein which exists as a dimer linked by a pair of disulfide bonds and exists in two types.^17^ The soluble form which is secreted by cells and the insoluble species which occurs in fibrillar extracellular structures.^17^ Fibronectin is ubiquitously expressed and involved in wound healing processes, cell adhesion, migration, growth and differentiation.^17^

Therefore, we investigated the expression of fibronectin during mouse and human kidney development in more detail and tested for its impact on nephrogenesis by deletion of fibronectin in *ex vivo* cultured metanephric mouse kidneys.

## Methods

### Human fetal kidneys

Formalin-fixed paraffin embedded sections of human fetal kidneys of week 10, 16, 21 and 35 of pregnancy were analyzed with permission provided by the local Ethics committee (reference number 4415). The causes for intrauterine death or abortion can only partially be provided. Three out of four cases were spontaneous abortions of unclear origin. In the fetus of week 35 of pregnancy fetal death occurred due to hemorrhagic shock of the mother with subsequent severe fetal anemia.

### Animals

Animal experiments were approved by the local institutional review board and all animal experiments complied with the United Kingdom Animals Act, 1986, and associated guidelines, EU Directive 2010/63/EU for animal experiments. Experiments were approved by the local Ethics Committee of the Government of Unterfranken/Wuerzburg (TS-2/2022 MedIV). Mice carrying the loxP-flanked conditional alleles of fibronectin (FN^fl/fl^) were kindly provided by Prof. Reinhard Fässler (Max Planck Institute of Biochemistry, Martinsried, Germany). Mice were crossed with CAGG-Cre-ER™ mice on a C57BL/6 background from Jacksons Laboratory that express Cre-recombinase after tamoxifen treatment in order to receive homozygous CAGG-Cre-ER™;FN^fl/fl^ mice. Cre-negative littermates were used as control for the fibronectin expression characterization of wild-type mice and non-tamoxifen induced mice served as control in the embryonic kidney culture experiments.

### Embryonic kidney culture and morphometric analysis

Metanephric kidneys were harvested from embryonic CAGG-Cre-ER™;FN^fl/fl^ mice at embryonic day E13.5. Therefore, embryonic kidney pairs from n=11 individual mice were cultured *ex vivo* on transparent Millicell organotypic cell culture inserts (Merck Millipore, Billerica, MA, USA) and maintained in a 37 °C humidified CO_2_ incubator for 5 days. For fibronectin deletion (FN^-/-^) one metanephric kidney was treated with (Z)-4-hydroxytamoxifen (HT; 500 nM, Sigma-Aldrich) whereas the contralateral kidney was treated with control medium only and served as control (FN^+/+^). HT was diluted in DMEM culture medium, containing 2 mM L-glutamine, 10 mM HEPES, 10 mM insulin, 5.5 μg/ml transferrin, 6.7 ng/ml sodium selenite, 32 pg/ml trijodthyronine, 250 U/ml penicillin, 250 µg/ml streptomycin, 25 ng/ml prostaglandin E and added below the culture inserts. Medium was changed after 24 h and 72 h. After 5 days, whole kidneys were fixed in paraformaldehyde (4%) and stained with FITC-conjugated dolichos biflorus agglutinin (DBA) to illustrate branching morphogenesis and for Wilms tumor protein (WT1) to depict glomerulogenesis. Metanephric kidneys stained with FITC-DBA and for WT1 were photographed along the z-axis providing a series of 20 photos covering the kidney from top to bottom. Obtained photos then were combined into a single extended depth of field image by the use of the “full focus” algorithm of the BZ-9000 analyzer software (V.2.1., Keyence, Japan). The resulting micrograph then was further processed by the use of ImageJ (NIH, V.1.45) comprising “subtraction of background”, “contrast enhancement”, “Filter mean” providing a blurred image and thereby reducing background noise, “conversion into 8-bit grayscale image”, “threshold”, “removal of outliers” which in some cases was extended by “manual excision” of outliers or artifacts always including the ureter. The resulting photo then was converted into a skeletal representation of the branching architecture by the use of “skeletonize” which then provided the number of branches by the use of the “analyze skeleton” algorithm. Since WT1 does not only stain for glomeruli but also for the cap mesenchyme, an additional algorithm was applied for the count of glomeruli by defining “circularity” values followed by the “particle analysis” algorithm that separates glomeruli from other (non-circular) structures as described previously.^18^ In addition, bright field depth of field images were captured along the z-axis to obtain kidney sizes. Kidney sizes, number of branches and glomeruli were compared to the contralateral control kidney.

### Immunohistochemistry and antibodies

Whole mount kidneys were stained with FITC-conjugated dolichos biflorus agglutinin (DBA; 1:500, Vector Laboratories, USA) and polyclonal rabbit anti-Wilms tumor protein (WT1; 1:500, abcam, Berlin, Germany) after 5 days of *ex vivo* culture. As secondary antibody anti-rabbit IgG AlexaFluor® antibody (1:1000; Thermo Fisher Scientific, Inc., Erlangen, Germany) was used. For analyses of fibronectin expression and localization at different time points during kidney development, mouse kidneys from E12.0, E13.5, E16.5, P0, P7 and adulthood comprising n=5-8 kidneys per time point and human kidneys from week 10, 16, 21 and 35 of pregnancy were used. Two-micron thick kidney sections were stained with polyclonal rabbit anti-fibronectin (1:2000, Dako, Denmark). Binding of the primary antibody was visualized with secondary donkey anti-rabbit antibody conjugated with AlexaFluor® 555 (1:1000, Molecular Probes, Invitrogen). UB branches and the collecting ducts were stained with FITC-conjugated DBA (1:500, Vector Laboratories, USA) and nuclei were visualized by the use of DAPI. Ki67 staining was performed using monoclonal rabbit anti-Ki67 antibody (1:100, Linaris, Dossenheim, Germany) and signals were amplified by the use of the Vectastain Elite ABC Kit (Vector Laboratories, Burlingame, CA) according to the manufacturer’s instructions. PCNA was stained by the use of mouse monoclonal anti-PCNA antibody (1:50; Dako) followed by the use of an ABC M.O.M. kit (Vector). ITGA8 was stained by the use of polyclonal goat anti-Integrin alpha 8 antibody (1:50; R&D Systems, Minneapolis) and GDNF was stained by the use of polyclonal rabbit anti-GDNF antibody (1:100; Bioss antibodies, USA). Signals were visualized using secondary donkey anti-rabbit antibody conjugated with AlexaFluor® 555 (1:1000, Molecular Probes, Invitrogen). Signals were analyzed with a DM6000B fluorescence microscope (Leica, Wetzlar, Germany), and photographs were taken with a Leica DFC 450C camera. Significance of co-localization was analyzed and visualized by the use of ImageJ (V.1.45) and the colocalization finder algorithm by Christophe Laummonerie and Jerome Mutterer (Institut de Biologie Moleculaire des Plantes, Strasbourg, France) as described previously.^19^ For quantification of Ki67, the color deconvolution algorithm (ImageJ) was applied to dissect the different signals from n=6 kidney pairs, followed by binarization and particle analysis to obtain the ratio of the number of positive cells and cortex area (normalized to mm^2^ cortex tissue). In addition, DBA-positive branches were dissected by marking them as regions of interest by the use of ImageJ and Ki67-positive cells were quantified within the marked branches and set in ratio to the total cell number. Therefore, a total number of n=145 branches from n=6 individual kidney pairs was analyzed. ITGA8, GDNF and WNT11 signals were quantified as described previously.^20^ Briefly, fluorescent signals from n=5-6 individual kidney pairs were turned into 8-bit images after subtracting background (ImageJ) and a predefined threshold was used for all images to capture the signals, which then were set in ratio to the whole tissue area. All analyses were performed in a blinded manner.

### Western blotting

Proteins were isolated from embryonic mouse kidneys using a sample buffer containing 50 mM Tris-HCl, 150 mM NaCl, 10 mM EDTA, 1% sodium deoxycholate, 0.1% SDS, 1% protease inhibitor mixture (Roche, cOmplete, EDTA-free, Mannheim, Germany) and 1% Triton X-100. Proteins were separated using NuPAGE 3–8% Tris-Acetate Protein Gels (Life Technologies/Gibco®, Karlsruhe, Germany). For the detection of fibronectin, proteins were blotted using an iBlot 2 Dry Blotting System (Thermo Fisher Scientific, Inc., Erlangen, Germany) to a polyvinylidene difluoride membrane (GE Healthcare Europe GmbH, Munich, Germany). Membrane was then incubated with primary anti-fibronectin (1:500; Dako, Santa Clara, USA) overnight. Proteins were visualized using horseradish peroxidase-conjugated secondary antibody and ECL detection. Monoclonal Anti-mouse Vinculin (1:4000, Novus Biologicals, USA) was used as loading control.

### Real-time PCR

RNA from embryonic kidneys was extracted by the use of the peqGold Total RNA-Kit (VWR International GmbH, Darmstadt, Germany) according to the manufacturer’s instructions. SYBR-Green-based real-time PCR was performed using StepOnePlus (Applied Biosystems, Foster City, CA, USA). Messenger RNA (mRNA) expression levels were normalized to 18S using the ΔΔCt method. All primer sequences are listed in Supplemental Table 1.

### ELISA

For quantitative detection of mouse GDNF protein release into the medium enzyme-linked immunosorbent assay (ELISA) was performed using the Mouse GDNF CLIA Kit (BIOZOL Diagnostica Vertrieb GmbH, Eching, Germany) according to the manufacturer’s instructions. For this analysis metanephric kidneys were harvested from embryonic CAGG-Cre-ER™;FN^fl/fl^ at E13.5 and cultured *ex vivo* on culture inserts for five days (n=6 kidney pairs). One of the kidneys was exposed to HT to induce fibronectin deletion (FN^-/-^) and the other kidney served as control (FN^+/+^). Medium was obtained after 24h, 72h and 120h and luminescence was measured using GloMax Microplate Reader (Promega GmbH, Walldorf).

### Statistical analysis

Data are expressed as mean ± SEM. An unpaired t test was applied to compare the differences between two groups; a paired t test was used for matched observations (kidney pairs). Wilcoxon signed-rank test for columns statistics was used for relative values. P < 0.05 was considered statistically significant.

## Results

### Fibronectin is expressed during kidney development with significant changes of its localization over time

First, we characterized the expression of fibronectin in wildtype mouse kidneys using immunofluorescent stainings at different time points of kidney development. Metanephric kidneys were harvested at E12.0, E13.5, E16.5, P0, P7 and adulthood. Fibronectin expression was detected during all stages of nephrogenesis. However, the localization of fibronectin signal changed significantly over time. At E12.0 and E13.5 fibronectin expression was primarily detected in the tubulo-interstitium, thereby strongly lining the ureteric bud branches (**Figure 1A, B**), which corresponded to previous studies.^16,21^ In embryonic kidneys at E16.5 fibronectin was still expressed in the tubulo-interstitium, but less pronounced around the ureteric bud epithelium, and extended by subtle expression within the (pre-) glomerular structures (**Figure 1C**). At P0 and P7 fibronectin expression disappeared from the tubulo-interstitium and changed to a strong glomerular staining pattern (**Figure 1D, E**). In healthy adult kidneys fibronectin appeared almost exclusively in the glomerular matrix (**Figure 1F**). Similar fibronectin expression patterns were found in human fetal kidneys at week 10, 16, 21 and 35 of pregnancy using immunofluorescence staining. In all those stages fibronectin expression was detected. Again, the pattern of fibronectin changed from tubulo-interstitial localization lining the tubular structures at week 10 of pregnancy (**Figure 1G**) to a more diffuse tubulo-interstitial expression at week 16 of pregnancy (**Figure 1H**). At week 21 and 35 of pregnancy fibronectin staining was predominantly present within glomerular structures (**Figure 1I, J**). These findings reveal that fibronectin is expressed during kidney development of mouse and human. The striking localization along the UB branches suggest that fibronectin may be involved in UB branching morphogenesis of developing kidneys.

**Figure 1.**
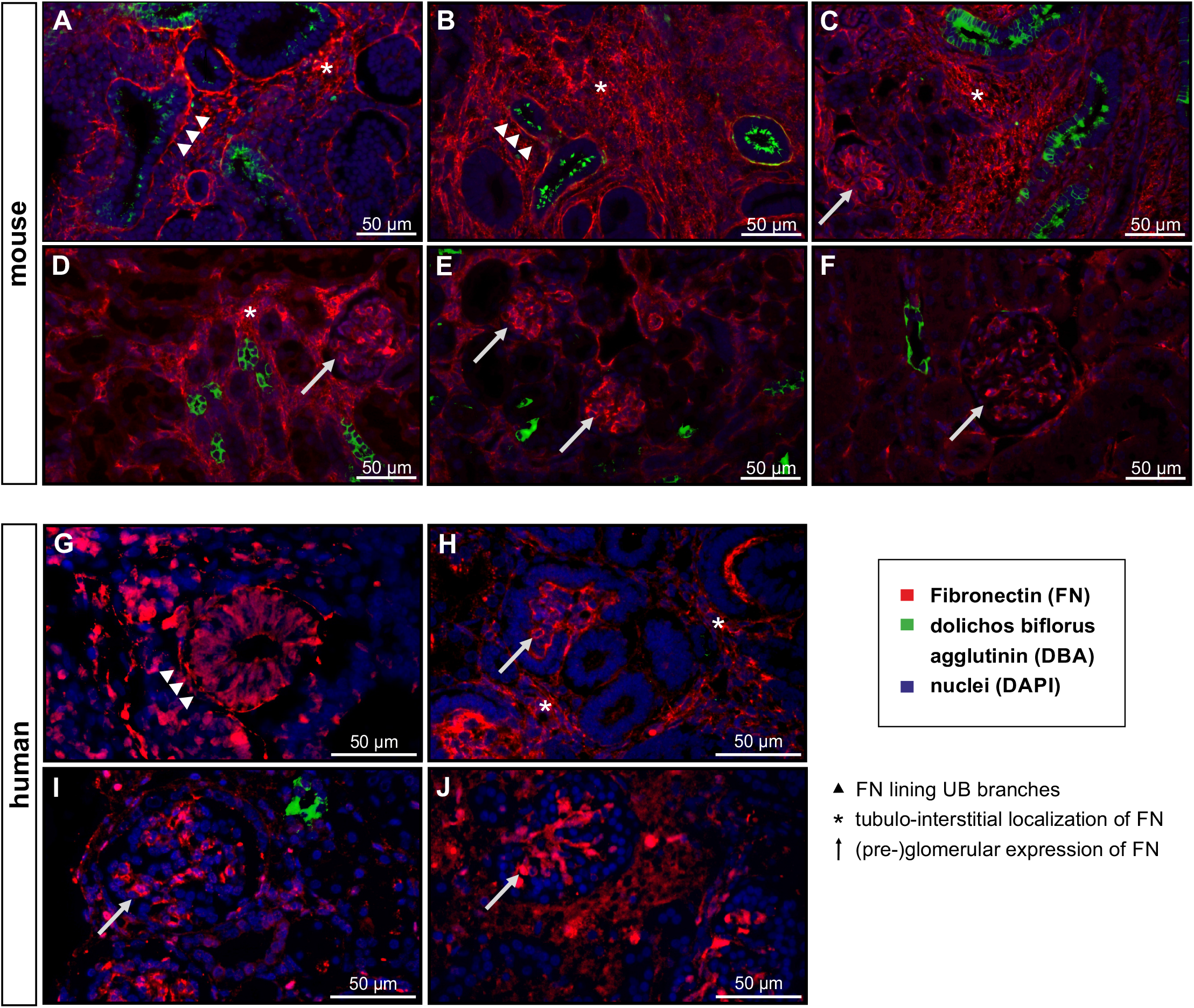
Expression and localization of fibronectin in kidney development. Fibronectin (red), dolichos biflorus agglutinin (DBA; green), as marker for UB cells and collecting ducts, and nuclei (DAPI; blue) were stained at different stages of kidney development. (**A-F**) show representative stainings of mouse kidneys at embryonic day E12.0 (**A**), E13.5 (**B**), E16.5 (**C**), postnatal day P0 (**D**), P7 (**E**), and adult kidney (**F**). (**G-J**) show representative stainings of fetal human kidney sections at week 10 (**G**), 16 (**H**), 21 (**I**) and 35 (**J**) week of pregnancy. Arrowheads indicate fibronectin lining ureteric bud epithelial cells, asterisks mark tubulo-interstitial fibronectin expression, and arrows indicate glomerular staining pattern.

### Tamoxifen-inducible deletion of fibronectin in ex vivo cultured metanephric kidneys

In order to study the role of fibronectin during embryonic kidney development CAGG-Cre-ER™; FN^fl/fl^ mice were generated allowing tamoxifen-inducible deletion of fibronectin. Metanephric kidneys were dissected at E13.5 and cultured *ex vivo* on organotypic Millicell organtypic cell culture inserts for five days. One kidney of the kidney pairs was genetically deleted for fibronectin by application of (Z)-4-hydroxytamoxifen (HT) (500nM) at day 1, whereas the contralateral kidney served as control (**Figure 2A**). Real-time PCR showed significant reduction of fibronectin mRNA level five days after application of HT compared to the contralateral control kidneys (**Figure 2B**). Additional Western blot analysis after five days of culture confirmed these findings on protein level (**Figure 2C**). Notably, significant reduction of fibronectin was already visible 48h after application of HT in kidney culture indicating a short half-life in kidney development (**Figure 2D)**.

**Figure 2.**
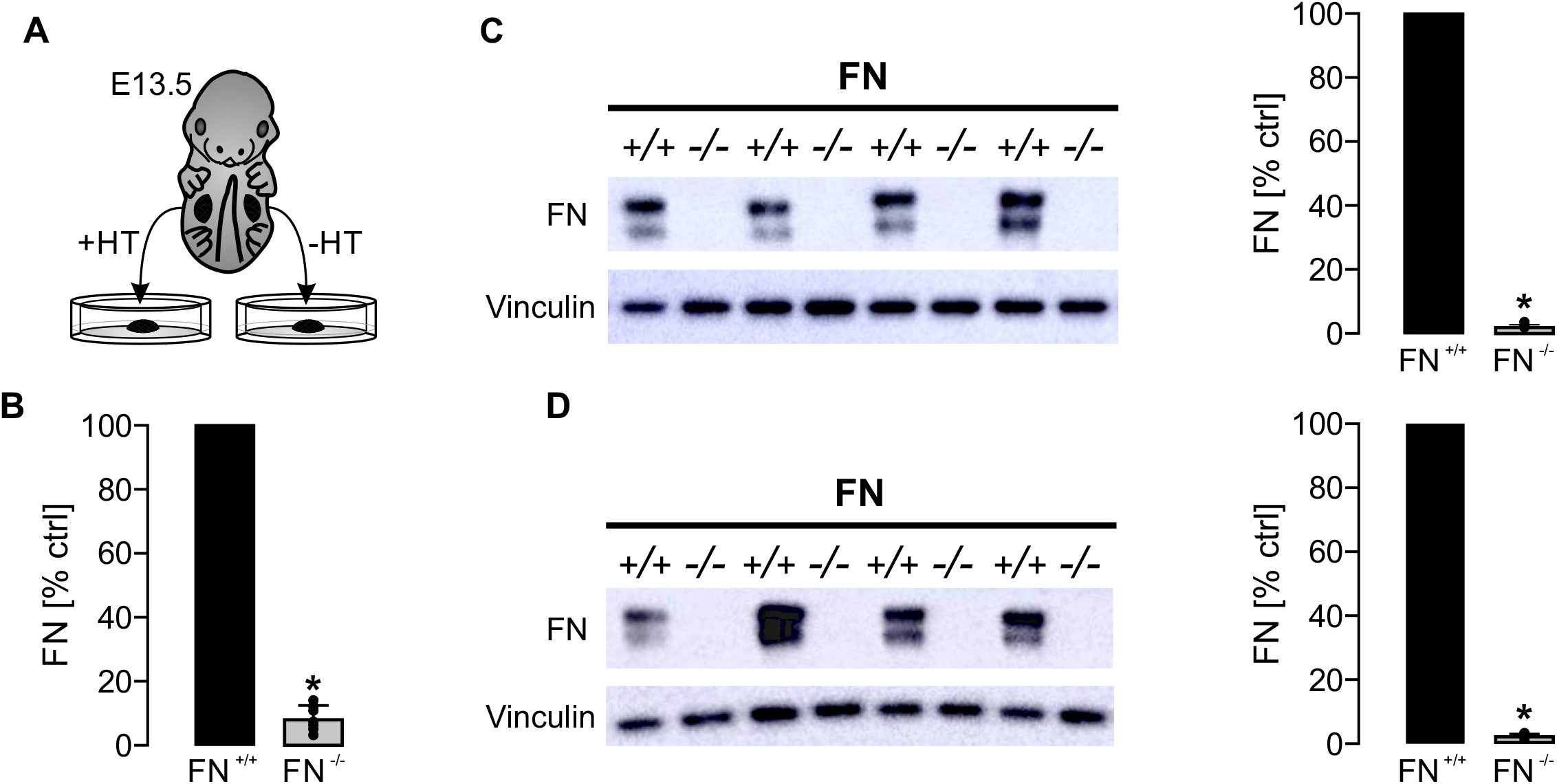
Fibronectin deletion in *ex vivo* cultured metanephric mouse kidneys. E13.5 metanephric kidney pairs (n=11) were cultured *ex vivo* for 5 days. (Z)-4-hydroxytamoxifen (HT) was applied after 24h of culture and the contralateral kidneys were maintained under control condition (illustrated in **A**). Application of HT resulted in a significant loss of fibronectin (FN^-/-^) 4 days later (day 5 of experiments) as indicated by Real-time PCR (**B**) and Western blot analysis (**C**). (**C**), left shows representative Western blot, right shows statistical analysis of fibronectin expression normalized for vinculin with FN^+/+^ set = 100%. +/+ represents control kidneys (not induced with HT), -/-represents contralateral kidneys that were induced with HT to delete fibronectin. (**D**) Significant reduction of fibronectin was already noticed 48h after application of HT by Western blot analysis (n=5 kidney pairs). *indicates significant reduction compared to FN^+/+^.

### Deletion of fibronectin resulted in reduced kidney sizes, impaired branching and lower number of glomeruli

Metanephric kidney pairs were dissected at E13.5 and cultured *ex vivo* for five days. One kidney was treated with HT to induce fibronectin deletion (FN^-/-^) whereas the contralateral kidney served as fibronectin-competent control (FN^+/+^). To test for the role of fibronectin in nephrogenesis whole mount kidneys were stained for UB branches and glomeruli and photographed at the end of day 5. Fibronectin-deleted kidneys were significantly smaller than control kidneys (**Figure 3A, D, G**). Furthermore, UB branching morphogenesis was significantly impaired upon deletion of fibronectin (**Figure 3B, E, H**). In addition, loss of fibronectin resulted in a reduced number of glomeruli (**Figure 3C, F, I**).

**Figure 3.**
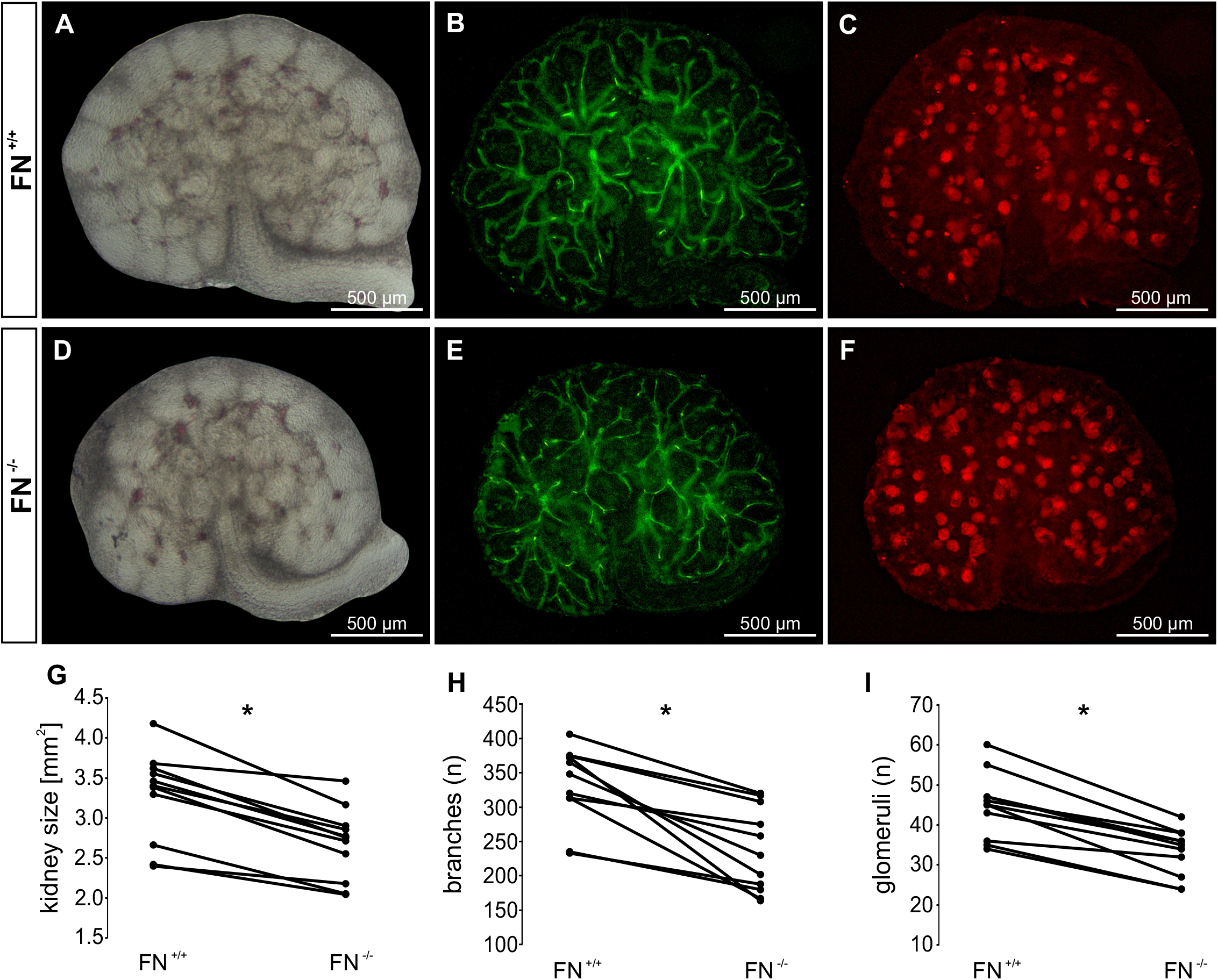
Loss of fibronectin resulted in reduced kidney size, impaired UB branching and lower number of glomeruli. E13.5 metanephric kidney pairs (n=11) were cultured *ex vivo* for 5 days. Deletion of fibronectin was induced at day 1 (FN^-/-^) and compared with contralateral kidneys cultured under control conditions (FN^+/+^) after 5 days of culture. Thereafter, kidney size, UB branches and glomeruli were visualized by the use of extended depth of field images of whole mount kidneys in combination with bright-field microscopy (**A, D**), DBA staining (green) (**B, E**), and staining for Wilms Tumor 1 (WT1; red) (**C, F**), respectively. (**G-I**) shows quantification of kidney sizes (**G**), number of branches per kidney (**H**), and number of glomeruli per kidney (**I**). * significant compared to FN^+/+^.

UB branching depends on cell proliferation.^22,23^ Therefore, kidneys were stained for the proliferation marker Ki67 five days after induction of fibronectin deletion. Ki67 signals were significantly reduced after deletion of fibronectin and normalized to the whole kidney tissue (**Figure 4A, B**). The effect was even more pronounced when Ki67-positive cells were analyzed exclusively within the UB branches and normalized to the total number of UB cells (**Figure 4A, C**). Similar results were obtained by the use of the proliferation marker PCNA (**Supplemental Figure 1**). These findings indicate that fibronectin promotes proliferation of UB branching cells which consequently affects kidney size and nephron endowment.

**Figure 4.**
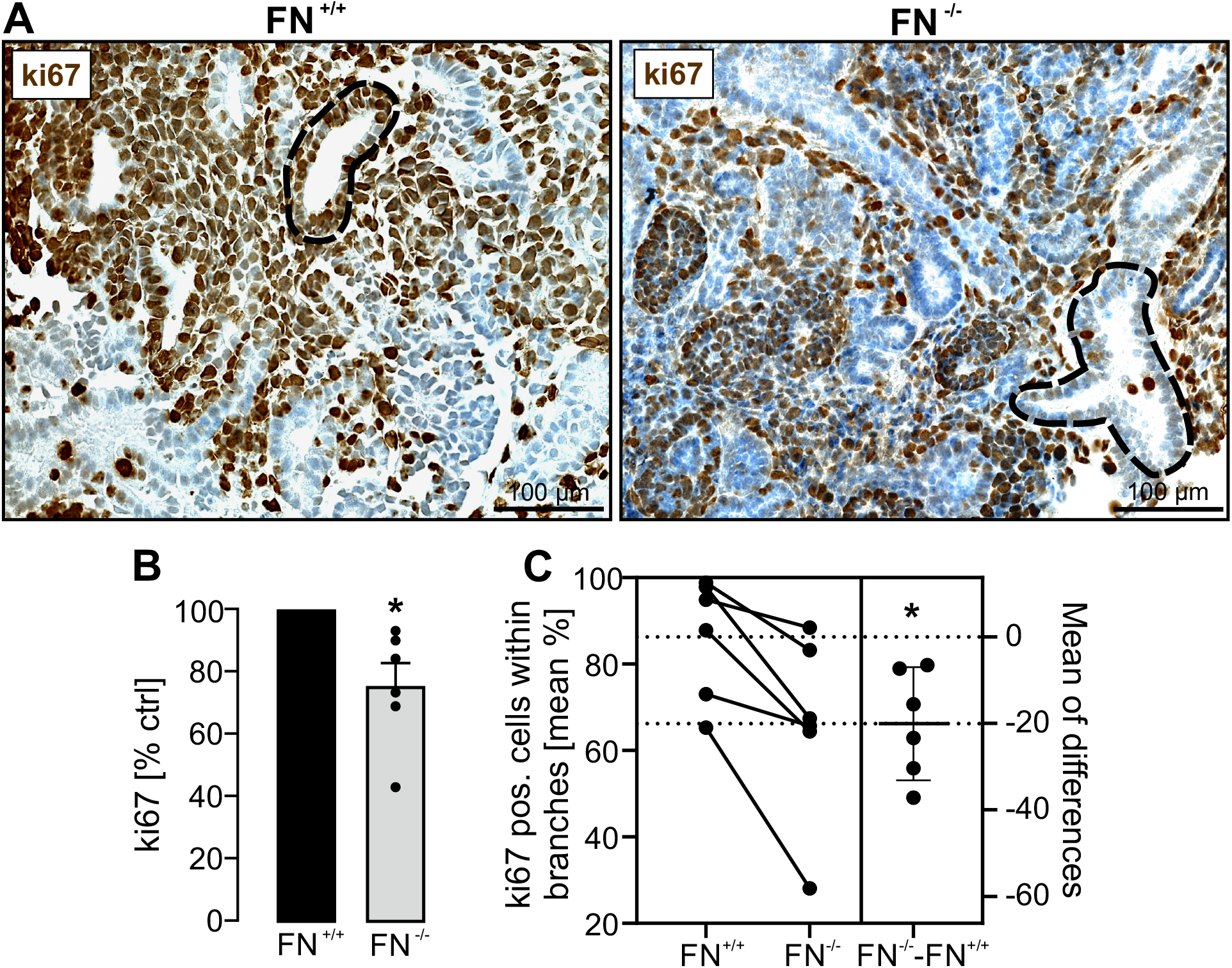
Loss of fibronectin resulted in reduced epithelial cell proliferation. Metanephric mouse kidneys (n=6 kidney pairs) were harvested at E13.5 and cultured *ex vivo* ± application of HT for five days. (**A**) Kidney sections were stained for the proliferation marker Ki67 (brown signals). The dotted line illustrates one UB branch in each of the photos. (**B**) Quantification of Ki67-positive cells in relation to total tissue area and normalized to FN^+/+^. (**C**) Quantification of Ki67-positive cells within UB branches and normalized to the total number of UB branching cells. * significant compared to FN^+/+^.

### Fibronectin colocalizes with ITGA8 in metanephric kidneys and affects ITGA8-dependent signaling

ITGA8 is involved in kidney development by promoting GDNF-mediated UB cell proliferation.^9,24^ Recently, the ECM protein nephronectin has been discovered as an important ligand to ITGA8 in kidney development.^15^ Fibronectin has also been suggested as a ligand to ITGA8, however with a much lower affinity.^25^ We next performed colocalization studies of fibronectin and ITGA8 and found significant colocalization of fibronectin and ITGA8 along the UB branches in metanephric mouse kidneys harvested at E13.5 (**Figure 5**). Surprisingly, deletion of fibronectin resulted in reduced expression of ITGA8 (**Figure 5**). We next stained for GDNF and found significant colocalization of fibronectin and GDNF in E13.5 metanephric mouse kidneys (**Figure 6A**). In line with the previous results, loss of fibronectin led to reduced expression of GDNF (**Figure 6B**) which was confirmed on mRNA level by the use of real-time qPCR (**Figure 6C**). In addition, we used ELISA assays to test for GDNF release into the medium over time after induction of fibronectin deletion compared to control kidneys (**Figure 6D**). Loss of fibronectin led to significantly reduced release of GDNF into the medium (**Figure 6D**). In line with these findings, real-time qPCR of cultured kidneys revealed a significant reduction of Wnt11 mRNA five days after induction of fibronectin deletion (**Figure 6E**).

**Figure 5.**
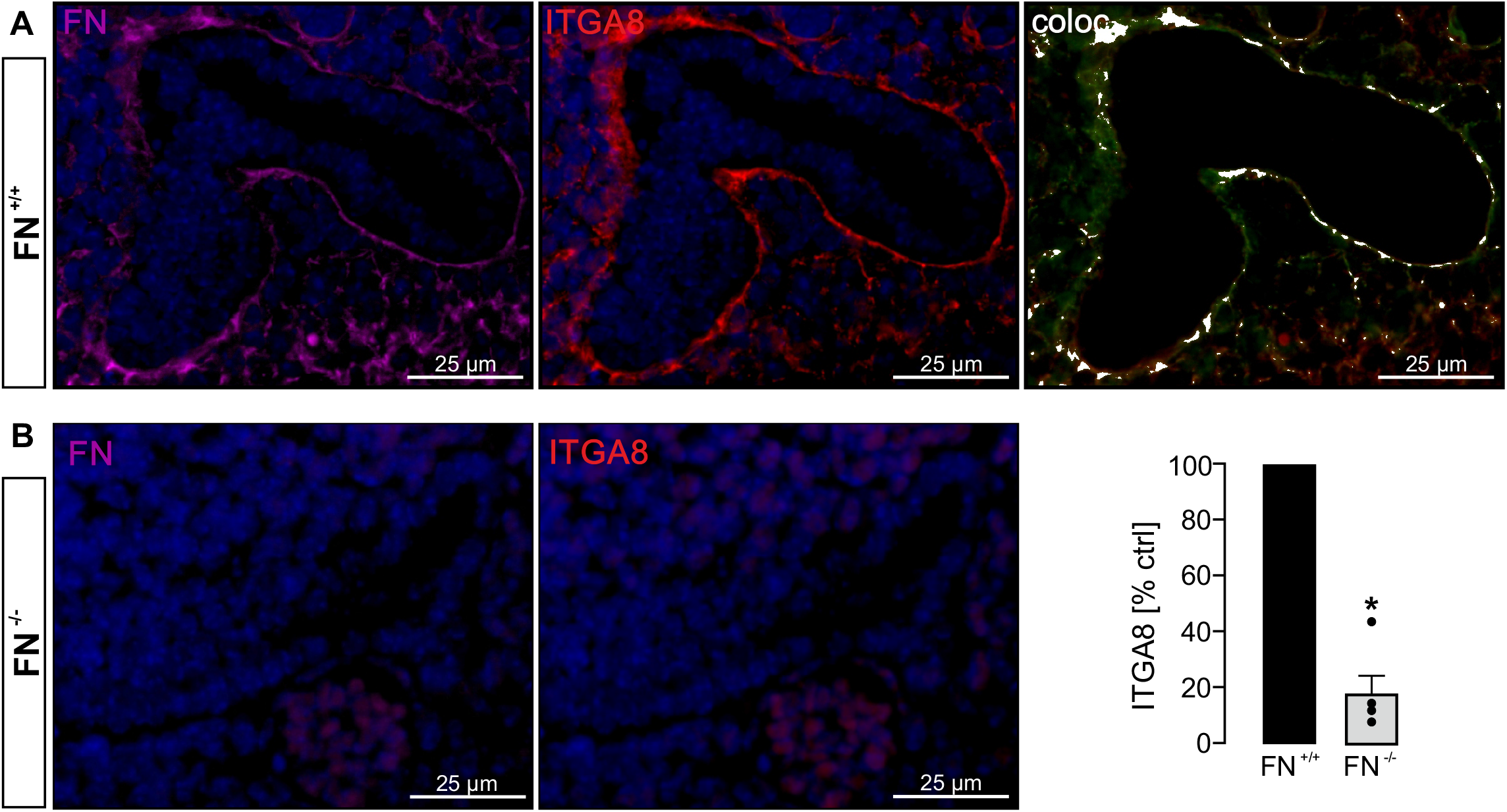
Fibronectin colocalized with ITGA8 and fibronectin deletion attenuated expression of ITGA8. Metanephric mouse kidneys (n=5 kidney pairs) were harvested at E13.5 and cultured *ex vivo* ± HT for five days. (**A**) Fibronectin-competent kidneys (FN^+/+^) were stained for fibronectin (FN; magenta), ITGA8 (red) and nuclei (DAPI; blue). Both, fibronectin and ITGA8 staining lined the UB branches and showed significant colocalization (white signals). (**B**) Fibronectin-deleted kidneys (FN^-/-^) showed reduced expression of fibronectin and ITGA8. Right shows quantification of ITGA8 staining signals in relation to total tissue area and normalized to FN^+/+^. * significant compared to FN^+/+^.

**Figure 6.**
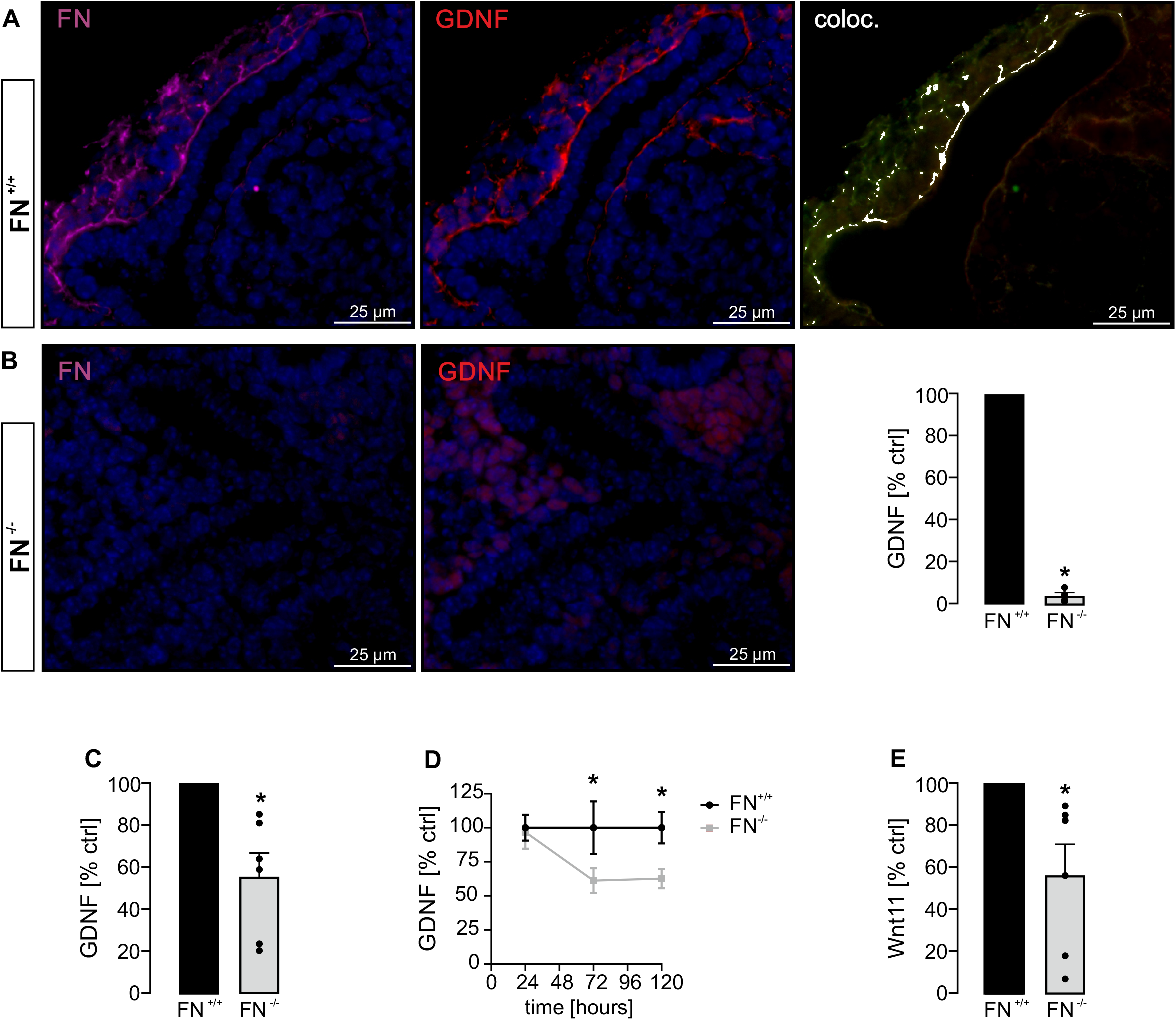
Fibronectin colocalized with GDNF and loss of fibronectin affected ITGA8-mediated signaling. Metanephric mouse kidneys (n=6 kidney pairs) were harvested at E13.5 and cultured ex vivo ± HT for five days. (**A**) Fibronectin-competent kidneys (FN^+/+^) were stained for fibronectin (FN; magenta), GDNF (red) and nuclei (DAPI; blue). Fibronectin and GDNF staining lined the UB branches and was significantly colocalized (white signals). (**B**) Loss of fibronectin resulted in lower expression of fibronectin and also of GDNF. Right shows quantification of ITGA staining signals in relation to total tissue area and normalized to FN^+/+^. (**C**) GDNF mRNA level was significantly reduced in fibronectin-deficient kidneys. (**D**) ELISA assays performed after 24h (time point of HT application), 72h and 120h revealed reduced concentration of GDNF in the medium of fibronectin-deleted kidneys. (**E**) Wnt11 mRNA expression was significantly reduced in kidneys upon deletion of fibronectin. * significant compared to FN^+/+^.

In summary, our data suggest that during kidney development fibronectin gets released by UB cells and stimulates ITGA8 in MM cells which leads to activation of GDNF. GDNF promotes UB cell proliferation and release of WNT11 (**Figure 7**).

**Figure 7.**
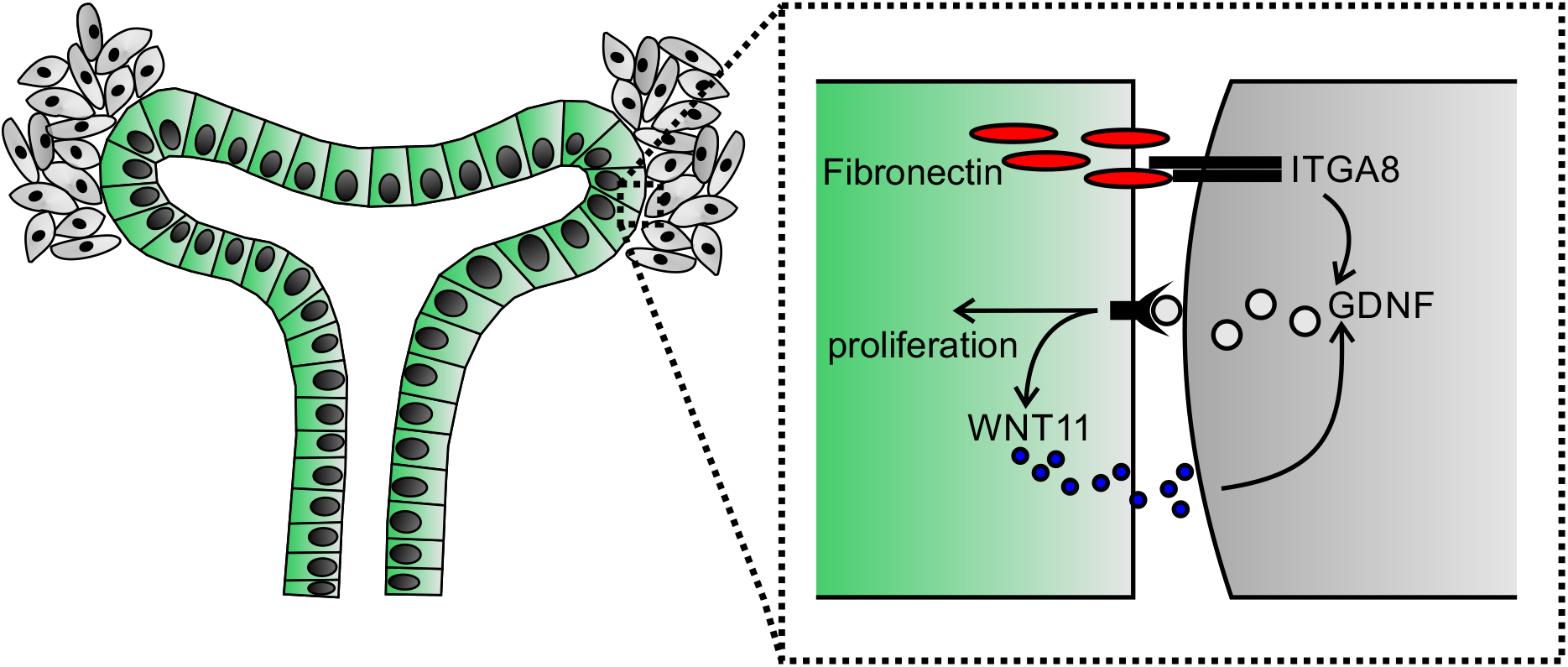
The involvement of fibronectin in kidney development. Our data indicate that during kidney development UB branching cells (green) release fibronectin which stimulates ITGA8 expressed by MM cells (gray). ITGA8 activation leads to release of GDNF which in turn promotes UB cell proliferation and release of WNT11. WNT11 further promotes GDNF release in a positive feedback loop.

## Discussion

Our study shows that the ECM protein fibronectin is expressed during kidney development and promotes UB branching. Therefore, fibronectin qualifies as a determinant of the final number of nephrons and glomeruli in the mature kidney.

Fibronectin is an ubiquitously expressed protein and involved in cell adhesion, growth, migration, and differentiation.^17^ Two types of fibronectin are present in vertebrates: soluble fibronectin, which is produced in the liver and representing a major protein component of blood plasma and insoluble cellular fibronectin, which is a major component of the ECM.^17^ The latter is secreted by various cells as a soluble protein dimer and later on is assembled into an insoluble matrix.^26^ Since we analyzed metanephric kidneys *ex vivo* and therefore isolated from blood flow, this indicates that endogenous fibronectin was responsible for the observed phenotype. The strong fibronectin signal lining the UB branches suggests the UB branching epithelial cells as the source of fibronectin with fibronectin being secreted from the UB cells as discussed by other groups before.^16,27^ In addition, similar expression pattern was observed in fetal kidneys. The human sample sizes were small and we cannot rule out that unknown underlying genetic defects or other alterations, which led to the spontaneous abortions may have affected fibronectin expression in these kidneys. However, given the high analogy to the mouse samples the findings indicate that the expression patterns may be representative.

Recently, the ECM protein nephronectin has been shown to promote UB branching by stimulation of ITGA8 and downstream-mediated release of GDNF.^15^ Knockout of nephronectin frequently resulted in renal agenesis^15^ whereas in our study deletion of fibronectin, which acts on the same signaling pathway, resulted in smaller kidneys with apparently intact architecture but reduced nephrons. There are several explanations for this discrepancy: Linton et al. used mice with a complete knockout of nephronectin^15^, which therefore may affect kidney development much earlier than in our model where we deleted fibronectin at E13.5. Induced deletion was necessary since complete lack of fibronectin leads to embryonic death in mouse before E10.5.^28^ Therefore, loss of fibronectin at an earlier stage may be more deleterious. In addition, induced deletion means that the protein has been generated until the time point of intervention and the following effects depend on the half-life of the protein. However, we show that already 48h after induction, fibronectin was largely reduced. Affinity of a ligand to its receptor is of course also noteworthy. Although fibronectin and nephronectin are both ligands of ITGA8, the affinity of fibronectin to ITGA8 is approximately 100-fold lower than that of nephronectin.^25^ This could mean that fibronectin may rather act as a modulator or fine tuner of ITGA8-dependent signaling. The latter idea would fit to a model where fibronectin may be a factor contributing to the determination of the final nephron number and not necessarily resulting in renal agenesis if its function is altered. Mutations in fibronectin have been linked to a particular renal disease, called glomerulopathy with fibronectin deposits (GFND, OMIM: 601894). GFND is a rare autosomal dominant disease characterized by proteinuria, microscopic hematuria, hypertension, and massive fibronectin deposits in the mesangium and subendothelial space, subsequently leading to end-stage renal failure.^29^ Recently, mutations localized in the integrin-binding domain in GFND patients have been detected.^30^ Although the disease is primarily regarded as being caused by massive fibronectin deposits it would be interesting (and challenging) to analyze if these patients also suffer from reduced nephron mass.

Low nephron number predisposes to arterial hypertension.^12,13^ Recently, in a genome-wide association study, a variant of fibronectin was linked to arterial hypertension.^31^ The underlying mechanism remained elusive. It is intriguing to speculate if this variant might impair kidney development and by this predispose to arterial hypertension. Given the ubiquitous expression and function of fibronectin, there are of course many other potential explanations for this observation.

In kidney research, ECM proteins are often regarded as acellular scaffolds that provide structural stability or as the unfortunate common histological end-point of progressive, chronic kidney diseases.^14^ Here, we show that the ECM protein fibronectin is involved in kidney development acting as a signaling molecule via ITGA8 receptors. Next to this novel physiological role, fibronectin variants with reduced or lacking binding affinity to ITGA8 may lead to lower nephron number and predispose to arterial hypertension and CKD. The latter is completely speculative at this point but may deserve attention in future research projects.

## Disclosure

All the authors declared no competing interests.

## Data sharing statement

Original data are available from the authors upon reasonable request.

## Acknowledgment

We thank Barbara Teschemacher, Andrea Luedke, and Eugenia Schefler for excellent technical support. BB and ET were supported by the Deutsche Forschungsgemeinschaft (DFG, German Research Foundation), project number 387509280, SFB 1350 (projects B3 and C2). This work was performed by KS in fulfillment of the requirements for obtaining the degree Dr. rer. nat.

## Supplementary Information

**Supplemental Table 1.**
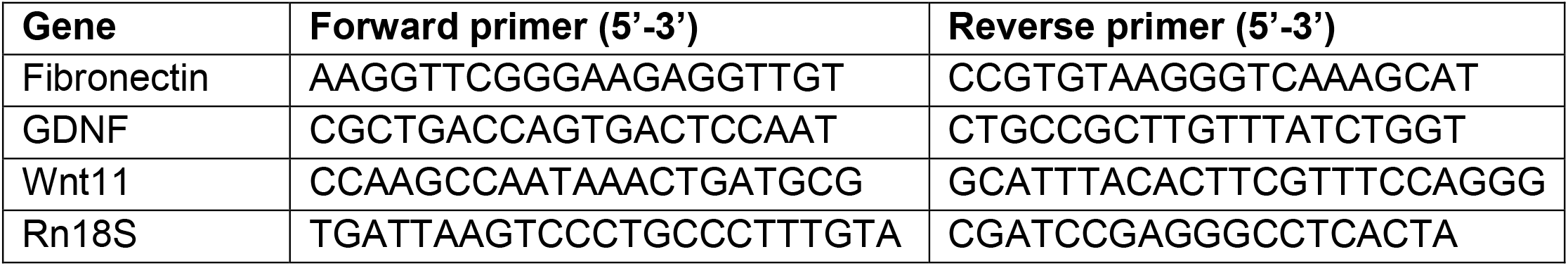
PCR primer sequences.

**Supplemental Figure 1.**
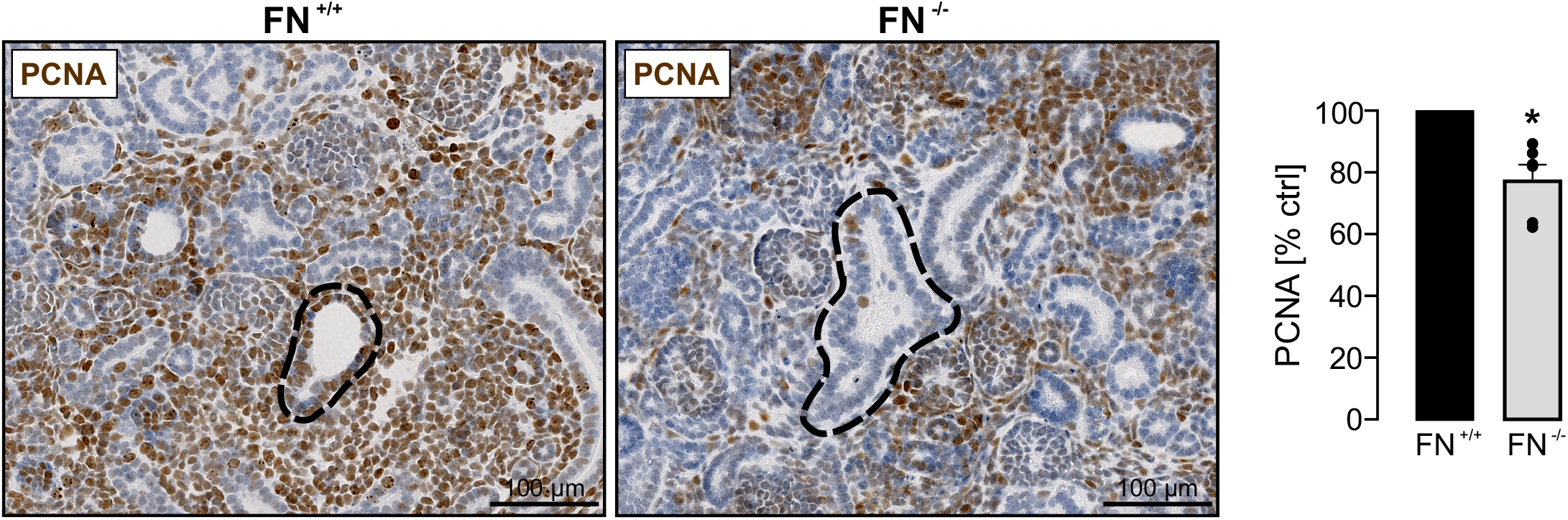
Loss of fibronectin resulted in reduced epithelial cell proliferation. Metanephric mouse kidneys (n=6 kidney pairs) were harvested at E13.5 and cultured *ex vivo* ± application of HT for five days. Left: Kidney sections were stained for the proliferation marker PCNA (brown signals). The dotted line illustrates one UB branch in each of the photos. Right: Quantification of PCNA-positive cells in relation to total tissue area and normalized to FN^+/+^. * significant compared to FN^+/+^.

## Notes

### Competing Interest Statement

The authors have declared no competing interest.

